# Negative frequency-dependent selection underlies overyielding through neighbor genotypic effects in *Arabidopsis thaliana*

**DOI:** 10.1101/2025.05.14.654149

**Authors:** Yasuhiro Sato

## Abstract

Negative frequency-dependent selection (FDS) can increase both genetic diversity and population-level mean fitness. While the dual consequence of negative FDS has long been theoretically recognized, its genomic basis remains unknown. To explore a genetic link between negative FDS and overyielding, I conducted a genome-wide association study (GWAS) of neighbor genotypic effects on the growth of 98 *Arabidopsis thaliana* genotypes. By incorporating neighbor genotypic similarity, my GWAS detected the most significant SNP on the fifth chromosome of *A. thaliana*, which was 6-kbp near a known locus responsible for indirect genetic effects. I found symmetric negative FDS on the most significant SNP, such that individual plants increased their biomass in the presence of neighbors with dissimilar alleles. I also observed overyielding in which a biallelic mixture of the most significant SNP showed a 3% increase in biomass compared to monoallelic conditions. Furthermore, I focused on the genomic region near the most significant SNP and found a signature of balancing selection near a negative regulator of ethylene responses *SIRTUIN2*. The present analysis uncovered a key locus linking balanced polymorphisms and overyielding in *A. thaliana*, showing a way to understand the maintenance of polymorphism and its positive impact on population productivity.

## 1. Introduction

Frequency-dependent selection (FDS) governs an important evolutionary regime that can decrease or increase genetic diversity within a population. It is widely recognized that negative or positive FDS maintains or unbalances polymorphism through rare-genotype advantage or disadvantage, respectively [1–3]. In addition to the maintenance (or not) of polymorphism, another important consequence of negative or positive FDS is an increase or decrease in population-level mean fitness [4–7]. This occurs as an increased or decreased fitness in polymorphic populations compared to additive expectations from monomorphic populations at a given locus [7], which may lead to overyielding [8]. In ecology, many researchers are aware of biodiversity effects wherein species and genotype mixtures become more productive than expected from monocultures [e.g., 9,10–12]. These relevant subjects in evolutionary biology and ecology highlight a potential connection between individual- and population-level productivity through the lens of FDS.

Despite being theoretically recognized, the impact of FDS on population-level mean fitness has rarely been demonstrated empirically [8,13,14]. A few pioneering studies reported negative FDS and a consequent increase in population-level mean fitness in insects [15,16]. In a damselfly *Ischnura elegans*, for example, rarer color morphs of females incur less mating conflict with males whereby negative FDS acts on the female color polymorphism [17]. Furthermore, a mixture of the two female morphs can increase egg production compared to monomorphic populations [15]. A reanalysis of the damselfly data [15] revealed that negative FDS underpinned egg overyielding in polymorphic over monomorphic populations [18]. Little is known, however, about how to identify such polymorphisms responsible for negative FDS and population productivity across a genome.

Genome-wide association study (GWAS) is increasingly recognized as a powerful tool to dissect the genetic architecture of adaptive traits in wild organisms [19–21]. Recently, Sato et al. [18] have proposed a method to enable GWAS of FDS on any fitness components. To detect FDS at each locus, this method employs regression analysis with allelic similarity considered a covariate. Based on the simplified FDS model, the covariate of allelic similarity represents negative or positive FDS based on its regression coefficient. Furthermore, a weighted mean between two alleles can be calculated as a result of estimated fitness functions, which represent population-level mean fitness in response to allele frequency. By repeating this regression for all single nucleotide polymorphisms (SNPs), we can conduct GWAS of FDS and thereby infer its potential influence on population-level mean fitness.

*Arabidopsis thaliana* is the best-studied plant species and thus its natural accessions provide an excellent opportunity to conduct GWAS for traits of ecological and evolutionary interest [22–24]. Based on the rich genomic resources of *A. thaliana*, recent studies have shown that overyielding occurs in a few combinations of focal and neighboring accessions [25,26]. Such indirect genetic effects from neighboring genotypes were shaped by ecological conditions and demographic history [27], which suggested that within-population genetic interactions were possible due to secondary contact between relict and non-relict lineages throughout Europe. These ecological and demographic processes are thought to exert balancing selection on genetic polymorphisms in *A. thaliana*, although a genetic link from neighbor effects via FDS to overyielding has yet to be explored.

To reveal the genetic link between FDS and population-level productivity, here I combined GWAS of FDS and pairwise cultivation data on natural accessions of *A. thaliana*. Because *A. thaliana* is monocarpic, plant size and biomass represent fecundity and resulting fitness [28]; thus, FDS on these fitness components was analyzed as follows. First, I narrowed down genomic regions associated with FDS by applying a recently developed GWAS method [18,29] to the pairwise cultivation data on *A. thaliana* [26]. I then compared patterns of plant biomass between biallelic and monoallelic conditions to examine whether negative FDS and overyielding co-occur at focal loci. Subsequently, candidate genes and genomic signatures of selection near the focal genomic region were sought to gain functional and evolutionary insights into the patterns of FDS and overyielding. Through the lens of FDS, the present analysis connects previous evidence of indirect genetic effects [27] and overyielding [26] with balancing selection.

## 2. Methods

### (a) Study species and data

The present study investigated the inbred and selfed genotypes called “accessions” of *A. thaliana* based on their open dataset. To analyze fitness components in *A. thaliana*, I reused phenotype data deposited by Wuest et al. [26] who quantified rosette size, flowering time, and individual biomass between 10 focal and 98 neighboring accessions. Wuest et al. [26] cultivated a pair of accessions in separated pots under a controlled greenhouse condition and deposited their data on Zenodo (see Data accessibility for DOI and URLs), which included 4172 individual plants in total. The genotype data for all the accessions were available from the RegMap project of *A. thaliana* [23] and its imputed SNP dataset with the 1001 Genomes project [30,31] (see Data accessibility for URLs). With the cut-off threshold of minor allele frequency (MAF) at 0.05, 206,416 and 1,897,419 SNPs were obtained from RegMap and imputed SNPs, respectively. As the 98 accessions belonged to the RegMap panel, the original RegMap variants and imputed SNPs were used for initial GWAS screening and post-GWAS analyses, respectively (see Data analysis below).

### (b) Model description

The present study adopted the regression models proposed by Sato et al. [18], which aimed to distinguish between negative and positive FDS through the effects of allelic similarity on a fitness component. In the present study, inbred *A. thaliana* accessions represented the specific case in which two homozygotes AA and aa interact within a population without mating over generations (see Appendix S2 “Case 3. Asexual or inbred lines without mating” of Sato et al. [18]). According to Sato et al. [18], I briefly describe this specific case as follows. Let *y*_*i*_ and *x*_*i*_ denote an observed fitness value and genotypic value of an individual *i* at a given locus, respectively. Then, let us assume that the focal individual *i* and *j* = 1,2, . . ., *N*_*k*_ individuals randomly interact within a subpopulation *k*, where *x*_*j*_ indicates a genotypic value of the counterpart individual *j* at the given locus. The following locus-level regression model formulates the effects of allelic similarity on a fitness component as:

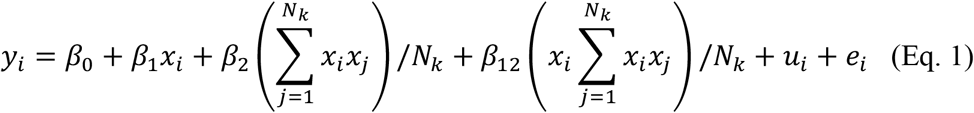

where the coefficients *β*_0_ and *β*_1_ represent the intercept and directional selection on *y*_*i*_, respectively. The other coefficients *β*_2_ and *β*_12_ respectively correspond to symmetric FDS and asymmetric FDS on *y*_*i*_ [18] when encoding two homozygotes as *x*_*i*_ ∈ {aa,AA} = {−1, +1}. This modeling is based on the Ising model of statistical physics [29] and thereby represents FDS when spatial structures are absent (see Eq. 4 below and Fig. 1). The random effect *u*_*i*_ and residual *e*_*i*_ represent variance components due to genetic and environmental factors, respectively. The random effect *u*_*i*_ considers three variance-covariance matrices such as directional selection, symmetric FDS, and asymmetric FDS (see Appendix S3 “mixed model extension” of Sato et al. [18]), which can correct for population structure due to these three factors [29,32]. When 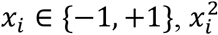 always turns 1 as 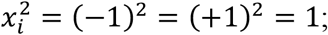 therefore, we can simplify the term of asymmetric 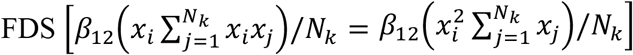 and thus exchange Eq. 1 as:

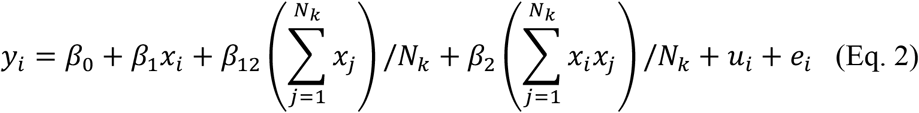

**Figure 1.**
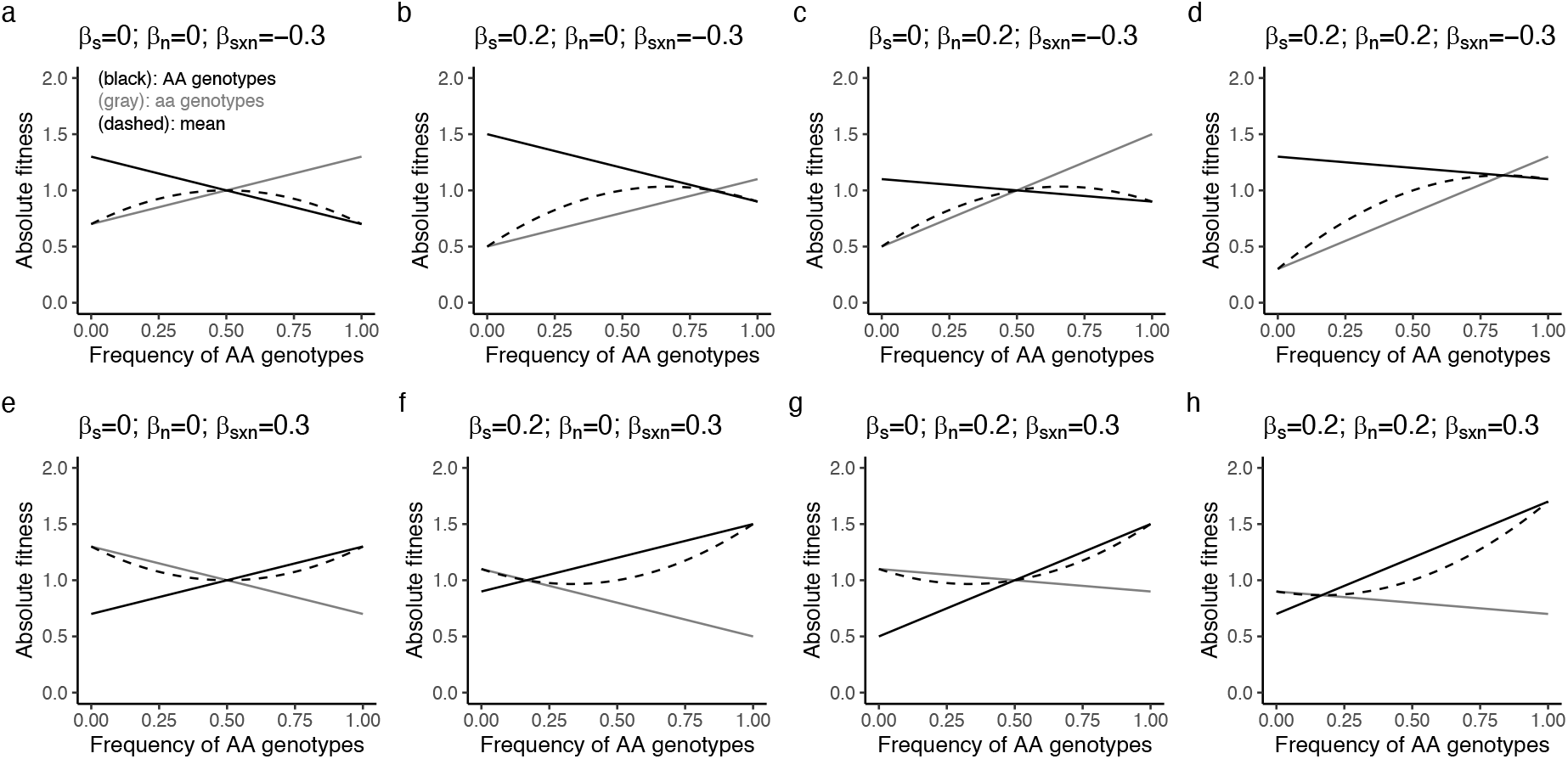
Numerical examples of frequency-dependent selection (FDS) and population-level mean fitness with respect to the regression coefficients of self-genotype effects *β*_s_, neighbor genotype effects *β*_n_, and self-by-neighbor genotypic effects *β*_s×n_. Y-axes show the absolute fitness value that corresponds to *y*_AA_, *y*_aa_, and 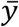 in Eq. 4. X-axes show the frequency of AA genotypes *f*_AA_ in Eq. 4. Black and gray lines indicate the fitness functions of AA and aa genotypes (*y*_AA_, *y*_aa_ in Eq. 4a and 4b), while dashed curves represent the population-level weighted mean between AA and aa genotypes (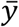 in Eq. 4c). (**a-d**) Examples of negative FDS on relative fitness, with modification of the absolute fitness conferred by *β*_s×n_ with modification by *β*_n_ and *β*_s_. (**e-h**) Examples of positive FDS on relative fitness conferred by *β*_s×n_ with modification by *β*_n_ and *β*_s_.

To distinguish the previous and present models, we redefine *β*_1_, *β*_12_, and *β*_2_ as self-genotype effects *β*_s_, neighbor genotypic effects *β*_n_, and self-by-neighbor genotypic effects *β*_s×n_, respectively. Then, Eq. 2 can be rewritten as:

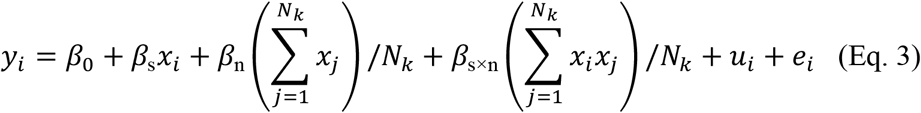

This Eq. 3 gives a multiplicative regression whose interaction term corresponds to symmetric FDS because of *x*_*i*_ ∈ {−1, +1}. Furthermore, when a subpopulation is so large that individuals freely interact with each other (i.e., *N*_*k*_ → ∞), Eq. 3 approximates *y*_*i*_ and accordingly presents fitness functions for two homozygotes and their weighted mean as:

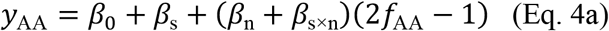

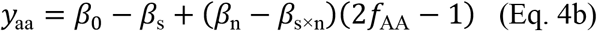

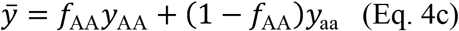

where fitness values of AA and aa homozygotes *y*_AA_ and *y*_aa_ are functions of AA genotype frequency (*f*_AA_). The population-level mean fitness 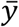 was then given by weighting *y*_AA_ and *y*_aa_ by their genotype frequencies. Based on Eq. 4a-c, Figure 1 shows numerical examples that represent negative/positive and symmetric/asymmetric FDS with its population-level weighted mean. For instance, when *β*_s×n_ < 0 and *β*_n_ = 0, symmetric negative FDS occurs with an increase in population-level mean fitness under biallelic mixtures. To link negative FDS and overyielding, Eq. 3 can be further simplified for the present case of interactions between a focal and neighboring individual as:

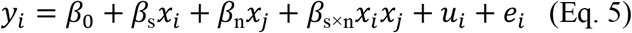

This specific case of two interacting individuals (Eq. 5) can address whether a given biallelic mixture increases or decreases *y*_*i*_ over monoallelic cultures through symmetric negative or positive FDS, respectively. Thus, I aimed to screen such SNPs with *β*_s×n_ < 0 and *β*_n_ = 0 in the following GWAS.

### (c) Data analysis

#### (i) Neighbor GWAS of plant fitness traits

Based on Eq. 5, I conducted GWAS of self, neighbor, and self-by-neighbor genotypic effects by testing the three coefficients *β*_s_, *β*_n_, and *β*_s×n_, respectively. Likelihood ratio tests for these three coefficients were performed from simpler to complex models following the procedure of Sato et al. [29]. I tested 206,416 SNPs from the RegMap panels individually to determine 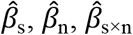 and their *p*-values. The target phenotypes were rosette size, flowering time, and individual biomass. These three phenotypes appeared to be normally distributed and thus were not transformed before GWAS. The community type (monoculture or mixture), experimental blocks, and plant position within a pot (A or B side) were included as non-genetic covariates. The gaston package [33] was used to solve the mixed models and perform GWAS, which followed the implementation of the rNeighborGWAS package [29]. All source codes generated by the present study are available through GitHub (see Data accessibility for URL).

To examine the significance of GWAS results, I diagnosed the inflation of *p*-values based on quantile-quantile plots and genome inflation factors [34]. When *p*-values were not inflated, Bonferroni correction was used to determine a genome-wide significance threshold at *p* = 0.05. When *p*-values were inflated, permutation tests were performed to empirically determine a significance threshold. Two types of permutations were used as the randomization of neighboring plants [32] or a genome-rotation scheme that shuffled the population genomic structure [35,36]. The empirical significance threshold at *p* = 0.05 was determined by 95 percentiles of the maximum -log_10_(*p*) association score per iteration among 199 iterations.

Once significant SNPs were detected by GWAS, I further screened SNPs with negative *β*_s×n_ to detect negative FDS and an increase in population-level mean fitness (i.e., the cases of Fig. 1a-d). To visualize changes in individual biomass, I plotted the observed individual biomass against biallelic or monoallelic conditions at focal SNPs. Linear regressions of biomass on biallelic or monoallelic conditions were performed for each allele to test whether genotype-level fitness functions were crossed within allele frequencies from 0 to 1. Welch’s *t*-test was also used to confirm biomass difference between the biallelic and monoallelic conditions, although the statistical significance of each SNP should follow that obtained from mixed models. The extent of biomass change was evaluated using a log-response ratio as log_e_(mean biomass under biallelic condition / mean biomass under monoallelic condition).

#### (ii) Fine mapping with imputed SNP data

To find candidate genes near the genomic region of focal SNPs, I conducted fine GWAS mapping using the imputed SNP data. The aforementioned GWAS was performed again using the full imputed SNP data between RegMap and the 1001 Genome project [30], which provided 1,897,419 SNPs for the 98 accessions. After mapping, LD and haplotype patterns near a significant SNP were examined to narrow down the location of candidate genes. The extent of LD was evaluated by the square of Pearson’s correlation coefficient *r*^2^ between two SNP loci among 98 accessions. Principal component analysis was used for the *r*^2^ correlation matrix to examine the genetic structure of focal genomic regions.

#### (iii) Genome scan of balancing and directional selection

A genome scan was performed to test whether the signatures of selection coincided with the genomic region of interest. Specifically, I analyzed balancing selection as the primary interest of this study and positive directional selection as a comparison. Both types of selection were analyzed based on local haplotype patterns using BETA [37] or EHH [38] method: The former and latter respectively detect allele frequency correlation and selective sweep at a haplotype level [37,38]. I used imputed SNP data with MAF cut-off at <0.05 for the 98 accessions. As described in Supplementary Note 5 of Sato et al. [32], I used multiple FASTA formats of the ancestral species *A. lyrata* genome to determine ancestral or derived alleles for each SNP. In total, I obtained 412,706 SNPs with known ancestral or derived allelic status. The genome-wide significance threshold was determined by 97.5 percentiles among all the SNPs for each type of selection, which corresponded to 5% significance in the context of two-sided tests.

To detect the signatures of balancing selection, I used Siewert and Voight’s BETA statistics that are based on haplotype-level nucleotide diversity around focal SNPs [37]. I used standard BETA statistics, that is, BETA^(1)^, because of the unknown divergence time between ancestral and derived alleles. The BETA scan was performed using a Python script available at the developer’s GitHub repository (see Data accessibility for URL), with default parameters adopted. This balancing selection index was called “BETA” throughout the present manuscript to avoid confusion with *β* coefficients of the regression models above.

To detect the signature of positive directional selection, I used the extended haplotype homozygosity and (EHH) its integrated haplotype score (iHS) [38]. The rehh package [39,40] in R was used to calculate EHH and iHS statistics. The default parameter settings were used in the scan_hh and ihh2ihs functions of the rehh package. I narrow down SNPs under balancing but not directional selection by comparing BETA and iHS statistics.

## 3. Results

### (a) Genomic region associated with negative FDS and overyielding

Among the nine GWASs of self, neighbor, and self-by-neighbor genotypic effects on biomass, flowering time, and rosette size (Fig. 2a-i), I detected many significant SNPs associated with the self-by-neighbor genotypic effects on biomass (Fig. 2g). While all GWASs were considered sound in terms of genomic inflation factors (≈ 1; Table S1), GWAS of self-by-neighbor genotypic effects on biomass displayed inflation of *p*-values in its QQ plot (Fig. S1g). I therefore performed permutation tests to determine a genome-wide empirical threshold for the biomass (Fig. S2). As a result, I found 21 significant SNPs on the first and fifth chromosomes beyond the empirical threshold of the significance level at *p* = 0.05 (above a grey horizontal line in Fig. 2g).

**Figure 2.**
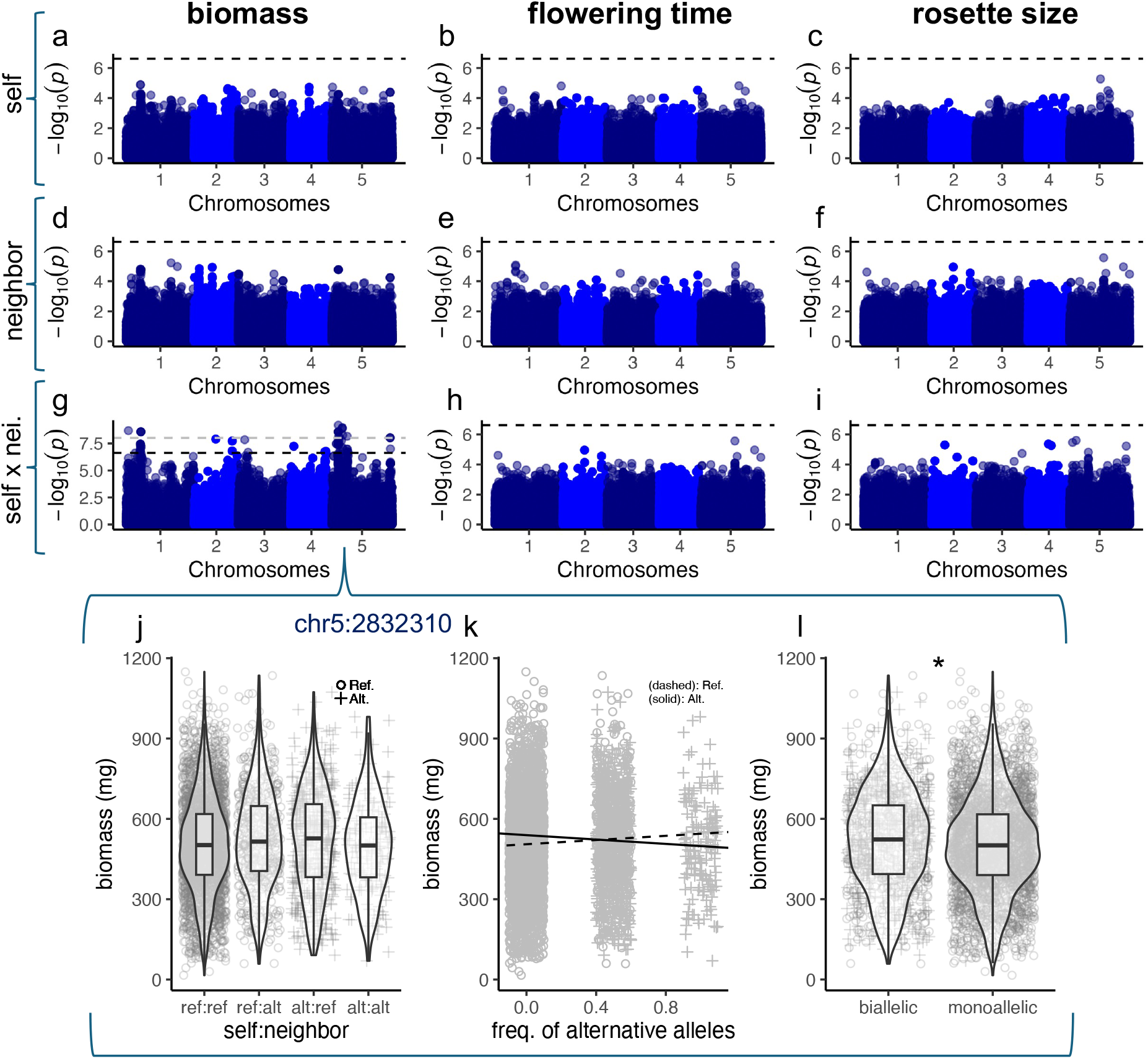
GWAS Manhattan plots for self (**a-c**), neighbor (**d-f**), and self-by-neighbor (**g-i**) effects on the biomass (**a, d**, and **g**), flowering time (**b, e**, and **h**), and rosette size (**c, f**, and **i**). Black horizontal dashed lines indicate the genome-wide Bonferroni threshold at *p* = 0.05. A grey horizontal dashed line in **g** indicates *p* = 0.05 by a permutation test (see Methods and Fig. S2a). The lower panels **j-l** focus on the most significant SNP at 2832310 on the fifth chromosome, where the biomass is plotted against self:neighbor allelic combinations between reference (ref) and alternative (alt) alleles (**j**); frequency of alternative alleles (**k**), and the biallelic or monoallelic conditions (**l**). Dashed and solid lines in (**k**) represent regression lines for individuals carrying reference and alternative alleles, respectively. In **l**, an asterisk (*) indicates a 5% significant difference between the biallelic or monoallelic conditions with a naive Welch’s *t*-test (*t* = 2.53, d.f. = 1674, *p* = 0.011). Box plots show the median with upper and lower quartiles, with whiskers extending to 1.5 × interquartile range.

I then focused on genomic regions near the 21 significant SNPs associated with self-by-neighbor genotypic effects on biomass. The most significant SNP on the fifth chromosome at 2832310 bp position exhibited patterns of negative FDS and overyielding (Fig. 2j-l). At this SNP locus, individual plants with rarer alleles produced more biomass than those with abundant alleles (Fig. 2k), resulting in a 3% biomass increase in the biallelic over monoallelic conditions (Fig. 2l). The set of significant SNPs on the fifth chromosome also included a SNP at 2838468 bp position, which was the same as previously detected SNP by GWAS of indirect genetic effects [27] (see also Discussion below). At this SNP (chr5:2838468), individuals with reference alleles produced more biomass with an increasing frequency of those with alternative alleles (Fig. S3d), but the two fitness functions did not cross to show patterns of negative FDS (Fig. S3e) and overyielding (Fig. S3f). Other than the fifth chromosome, significant SNPs were also detected on the first chromosome (Fig. 2g), although the most significant SNP on the first chromosome did not exhibit patterns of negative FDS and overyielding (Fig. S3a-c). Taken together, these results showed that the most significant SNP on the fifth chromosome exhibited a pattern of negative FDS and overyielding, leading me to focus on this genomic region.

### (b) Quantitative trait locus resolved by fine mapping

To resolve the genomic region on the fifth chromosome and find candidate genes therein, I performed GWAS using the higher-resolution SNP data [30]. Among SNPs on the fifth chromosome, this fine-mapping identified the most significant SNP at 2832310 bp position in addition to other SNPs with similar -log_10_(*p*) association scores (Fig. 3a), forming a quantitative trait locus. This locus spanned an approximately 100 kbp region that included 34 candidate genes (Fig. 3a and 3b). To further examine the genetic structure of this 100 kbp genomic region, I performed a principal component analysis and found that the first to fourth principal components explained 94% of the genomic variation (Fig. S4c). The patterns of haplotype and LD were consistent with the results of the principal component analysis, which showed approximately four LD blocks (Fig. S4a and S4b). One of the four haplotype blocks encompassed the aforementioned SNPs responsible for indirect genetic effects (i.e., chr5:2838468 [27]) and negative FDS (chr5:2832310) despite their weak LD (*r*^2^ = 0.017), indicating a potential link between the previous and present SNPs. In total, further dissection of the significant genomic region narrowed down the candidate genes and population genetic structure.

**Figure 3.**
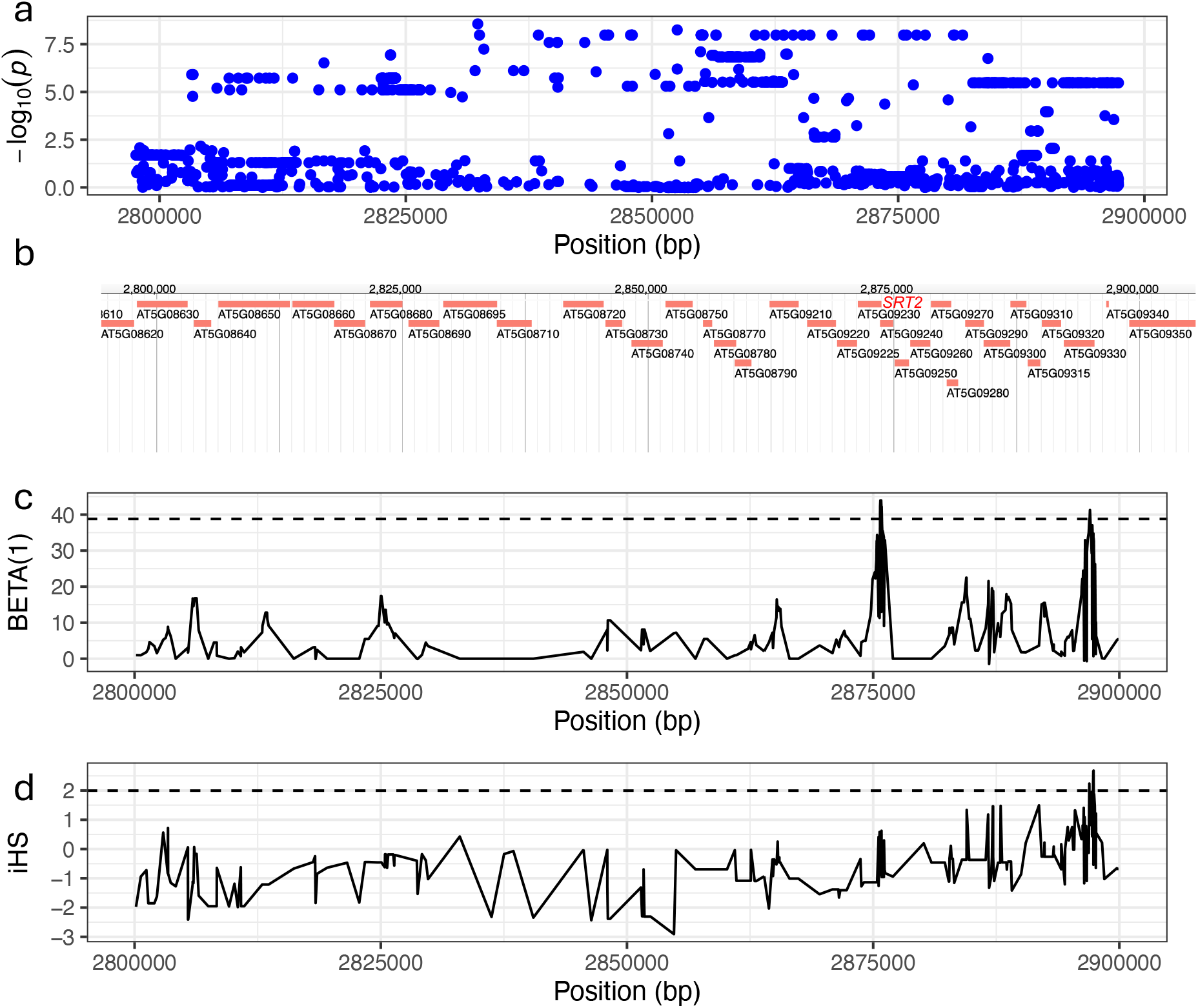
Fine mapping and signatures of selection near the most significant SNP on the fifth chromosome. (**a**) Fine mapping based on the imputed SNP dataset of *Arabidopsis thaliana*. (**b**) Candidate genes located near the focal genomic region. (**c**) Signatures of balancing selection based on BETA statistics (but different from *β* in Figure 1) of allele frequency correlation. (**d**) Signatures of directional positive selection based on the integrated haplotype score (iHS) of extended haplotype homozygosity (EHH). Horizontal dashed lines indicate genome-wide 97.5 percentiles of the two selection statistics.

### (c) Genomic signatures of balancing and directional selection

Finally, I conducted a genome scan of balancing and directional selection to narrow down candidate genes with the signature of balancing rather than directional selection (Fig. S5). Within the focal region of the fifth chromosome, two regions showed relatively strong signatures of balancing selection BETA1 = 43.93 and 41.26 near SNPs chr5:2875787 and chr5:2897015, respectively) (Fig. 3c), which ranked higher than the 97.5 percentile in the genome-wide BETA distribution (Fig. S5a and S5b). One of these two regions also had a signature of directional selection (Fig. 3d), whereas the other region underwent balancing selection alone (Fig. 3c and 3d). I therefore focused on the latter region and found the *SIRTUIN 2* (*SRT2*) locus (near chr5:2875787 in Fig. 3c), which encodes a histone deacetylase, is required for the negative regulation of ethylene-responsive genes, and is involved in root length regulation when treated with 1-aminocyclopropane-1-carboxylic acid [41]. The joint evidence from phenotype-based GWAS and phenotype-free genome scan suggests that balanced polymorphisms can underpin negative FDS and overyielding, with functional insights provided by candidate genes (see also Discussion below).

## 4. Discussion

The present reanalysis of pairwise cultivation data identified the most significant SNP associated with negative FDS and overyielding on the fifth chromosome of *A. thaliana*. The nearby genomic region of this SNP also included the other significant SNPs and thus formed a quantitative trait locus associated with FDS. These results could be directly compared to those of previous studies on overyielding [25,26] and indirect genetic effects [27] because these present and previous studies were based on the same dataset. Notably, the present analysis detected the two significant SNPs at 2832310 and 2838468 bp positions within the same haplotype block, the latter of which was the same SNP as one of 11 significant SNPs responsible for indirect genetic effects on biomass [27]. On the surrounding 100 kbp region of the two SNPs at 2832310 and 2838468 bp positions, Montazeaud et al. [27] detected the signature of past admixture and introgression between relict and non-relict *A. thaliana* accessions [27]. This previous result suggested that indirect genetic effects were shaped by past demographic events in *A. thaliana* [27]. In addition to the previous evidence, the present findings provide further evidence that past demographic events might have contributed to shaping negative FDS as well as indirect genetic effects in *A. thaliana*.

Besides the demographic pattern, Montazeaud et al. [27] detected a significant genome-environmental association for soil water availability near their significant SNP on the fifth chromosome at 2838468 bp position, suggesting the relevance of belowground interactions to their indirect genetic effects in *A. thaliana*. In the present study, the significant genomic region on the fifth chromosome spanned a broad range that encompassed 34 candidate genes (Fig. 3b). These multiple candidates ultimately need to be validated using mutant lines, but would add functional insights to negative FDS and overyielding. Specifically, the genomic signatures of balancing selection led us to detect *SRT2* as a candidate gene that could be involved in root functional traits and belowground interactions. Regarding belowground interactions, Wuest et al. [25] found that allelic variation at the *AtSUC8* locus, which is located on the third chromosome, conferred root plasticity and consequent overyielding [25]. Interestingly, they also found that root plasticity was induced under acidic belowground conditions [25], while the present candidate gene *SRT2* is also known to play a role in acid-induced root elongation [41]. Although a direct link and experimental proof are needed, the present results provide supportive evidence for the previously reported roles of belowground interactions and root development in mediating indirect genetic effects and overyielding.

Theoretical studies suggest that protected polymorphisms and population-level mean fitness could be regarded as two sides of the same coin, such that symmetric negative FDS can increase population-level mean fitness at an intermediate frequency [5–7]. Although this dual consequence of negative FDS has been demonstrated in insects [15,16], there is limited evidence for plants. This may be due to the difficulty in distinguishing directional selection and FDS on traits relevant to environmental adaptation in sessile organisms [18,42]. The present method overcame this issue by decomposing the effects of neighboring genotypes into locus-wise FDS to enable the genome-wide screening of SNPs associated with FDS and overyielding. Although 3% overyielding may not appear large, this percentage is plausible compared with quantitative genetics and GWAS of indirect genetic effects. A recent meta-analysis has documented that the phenotypic variation explained by indirect genetic effects was 3% on average [43]. At the SNP level in GWAS, Montazeaud et al. [27] reported that joint indirect genetic effects from the top 11 SNPs explained 2.3% of the total phenotypic variation in *A. thaliana* biomass. The present GWAS of the model plant species showed values consistent with those of previous studies, providing evidence for the quantitative control of negative FDS and overyielding.

While the present study found negative FDS and overyielding, positive FDS and decreased population-level mean fitness could also be detected. These opposite outcomes can be distinguished using the present method, because self-by-neighbor genotypic effects *β*_s×n_ determine the concave or convex pattern of population-level mean fitness along genotype frequencies irrespective of self and neighbor genotypic effects *β*_s_ and *β*_n_ (Fig. 1). This implication aligns with the previous theory of FDS and biodiversity effects, in which negative or positive FDS increases or decreases net biodiversity effects, respectively [16]. Takahashi et al. [16] further partitioned net biodiversity effects into selection and complementarity effects using Loreau and Hector’s method [44], where negative FDS can enhance complementarity effects on population-level mean fitness. Based on the three parameters *β*_s_, *β*_n_, and *β*_s×n_, the present method may be more generally applicable than the case of symmetric negative FDS to understand how intraspecific biodiversity effects occur under FDS.

Other than biological concerns, modeling issues should be noted when applying multi-factor mixed models for GWAS. To correct the population structure more accurately, two-way mixed models of indirect genetic effects have often considered covariance between the self and neighbor genotypic effects [27,45]. However, the inclusion of covariance requires the full computation of mixed models for all SNPs due to the non-additivity of multiple variance components and thus makes GWAS difficult. Instead of including covariance, an interaction term between self and neighbor genotypic effects was incorporated as a non-additive component in the present locus-wise FDS model. This enabled the use of an additively weighted kinship matrix for diagonalization as implemented in many GWAS practices [33,46–48]. Despite the model extension, we still faced the issue of inflated *p*-values of self-by-neighbor genotypic effects on biomass. While the issue of inflated *p*-values has been empirically addressed by permutation tests, this approach is computationally demanding and not widely applicable to GWAS. Not only genomics but also theoretical investigation of advanced mixed models is needed for efficient implementation of GWAS of locus-wise FDS.

In conclusion, the present GWAS uncovered a key locus linking negative FDS and overyielding in a plant species. While the present study focused on growth traits, the same analysis can be applied to a wide variety of traits such as pathogen and herbivore resistance [32,49]. Having applied the same methodology to field-grown *A. thaliana*, Sato et al. [32] indeed identified key genotype pairs that could mitigate herbivore damage in genotype mixtures over monocultures. Maximization of productivity and resistance by genetic diversity provides a potential solution to reconciling biodiversity and ecosystem services in nature and agriculture [50,51]. To this end, modern statistical tools and genomic resources will enable a quantitative test for the consequences of FDS on both the maintenance and function of genetic diversity.

## Ethics

This work did not require ethical approval from a human subject or animal welfare committee.

## Data accessibility

All source codes generated by this work are available at the GitHub repository (https://github.com/yassato/Ara98biomass). Phenotype data are deposited by Wuest et al. [26] and available at Zenodo (https://doi.org/10.5281/zenodo.2659734). Genotype data for RegMap and imputed SNPs are available at the website of Joy Bergelson Lab (https://bergelsonlab.org/resources/a-thaliana/#regmap; accessed on 27-Nov-2024) and AraGWAS Catalog (https://aragwas.1001genomes.org/#/download-center; accessed on 27-Nov-2024), respectively. The Python script of BETA scan was downloaded from GitHub (https://github.com/ksiewert/BetaScan; accessed on 27-Nov-2024).

## Declaration of AI use

I have not used AI-assisted technologies in generating this article.

## Authors’ contributions

Y.S.: conceptualization, data curation, formal analysis, methodology, visualization, funding acquisition, project administration, writing - original draft, writing - review and editing.

## Acknowledgements

I am grateful to Samuel E. Wuest for invaluable feedback and comments on the manuscript. The super-computing resource was provided by Human Genome Center at the University of Tokyo (http://sc.hgc.jp/shirokane.html).

## Supplementary materials

**Figure S1.**
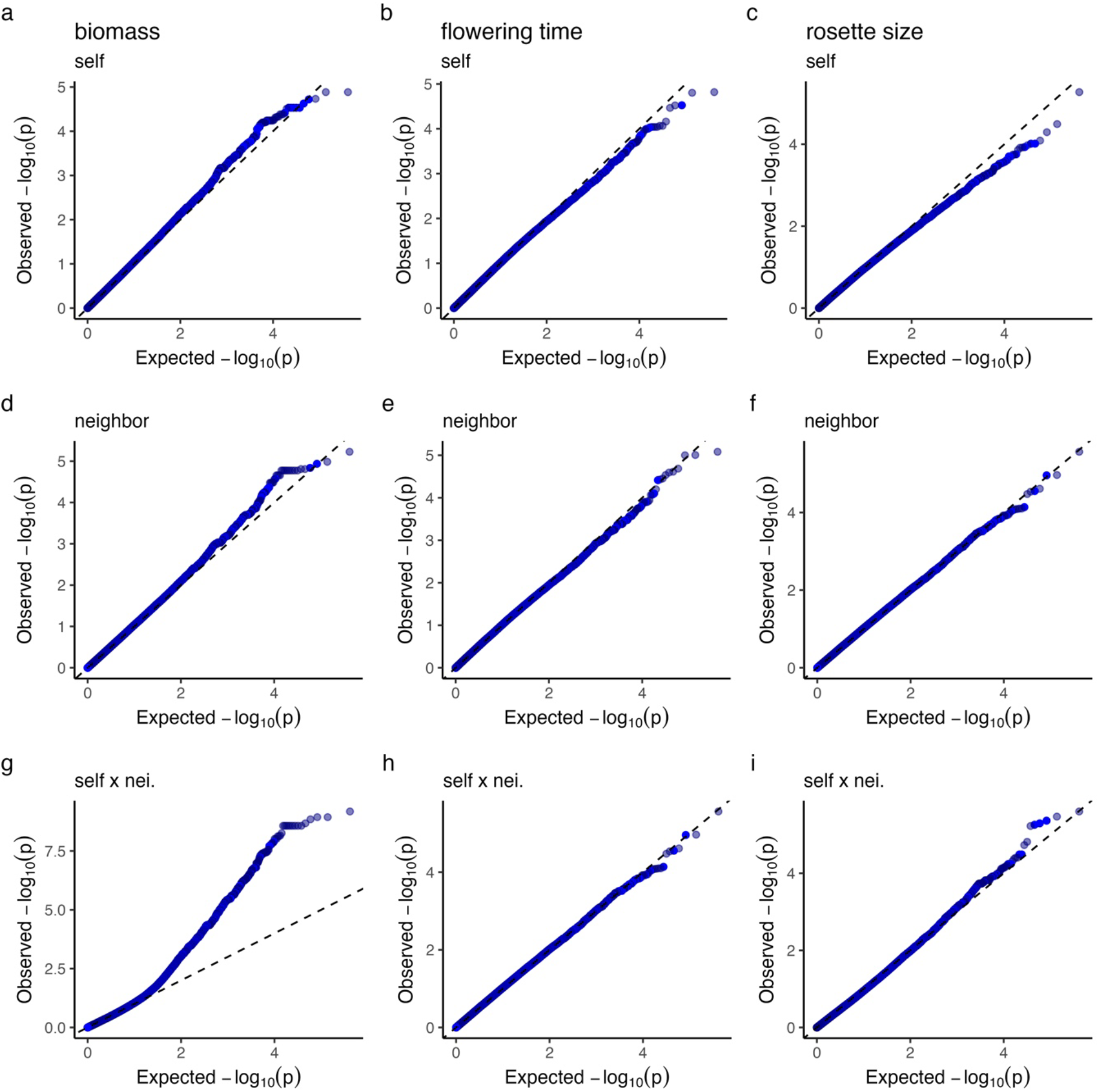
Quantile-quantile (QQ) plots of *p*-values for self (**a-c**), neighbor (**d-f**), and self-by-neighbor (**g-i**) effects on the biomass (**a, d**, and **g**), flowering time (**b, e**, and **h**), and rosette size (**c, f**, and **i**). X- and Y-axes show the expected and observed -log_10_(*p*), respectively. Dashed lines indicate *y* = *x*.

**Figure S2.**
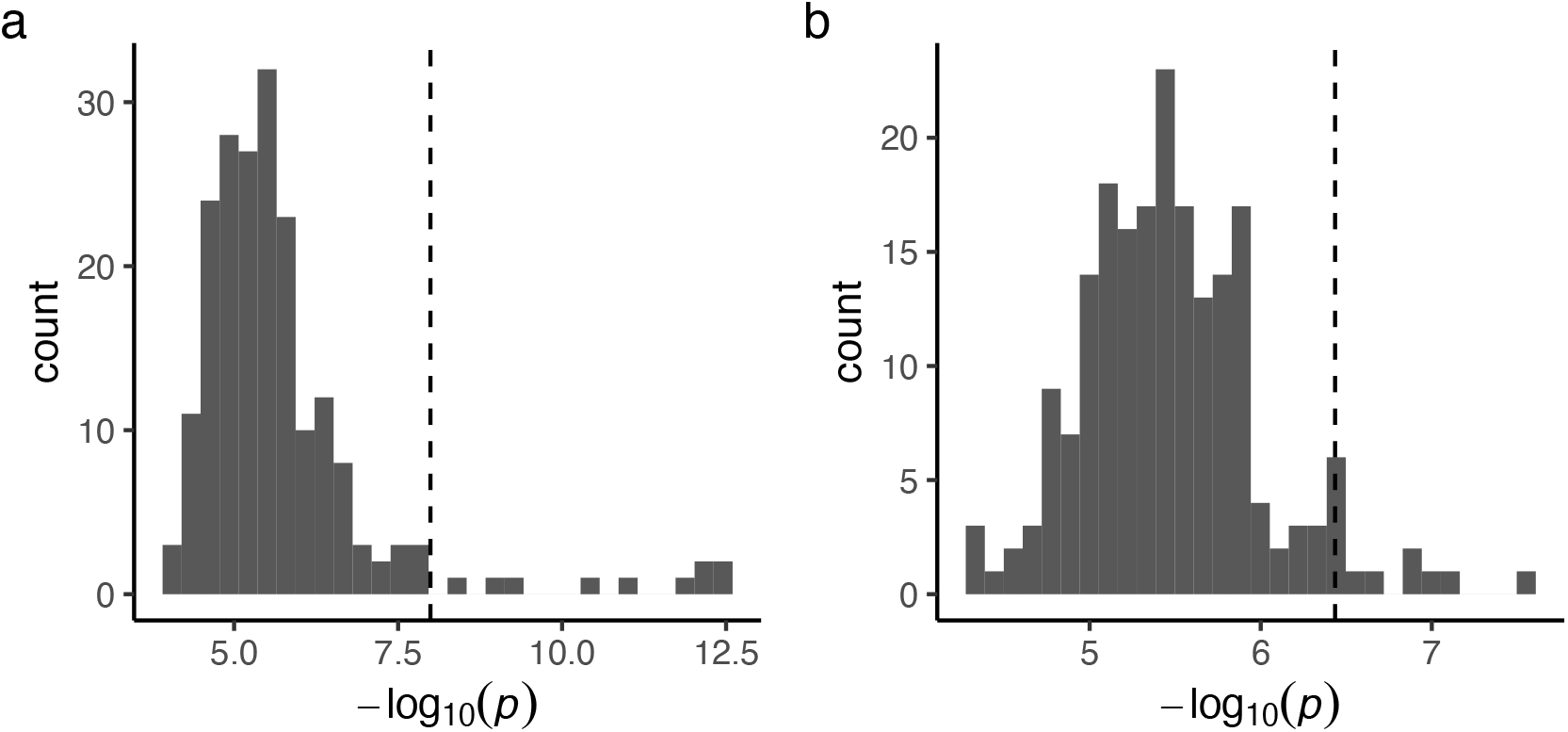
Distributions of the maximum -log_10_(*p*) among 199 permutations based on (**a**) random shuffling of neighboring plants or (**b**) genome-rotation scheme for self-by-neighbor GWAS *p*-values of biomass. Vertical dashed lines indicate 95 percentiles of the maximum - log_10_(*p*). Note that a more conservative threshold (**a**) was applied to Figure 2g.

**Figure S3.**
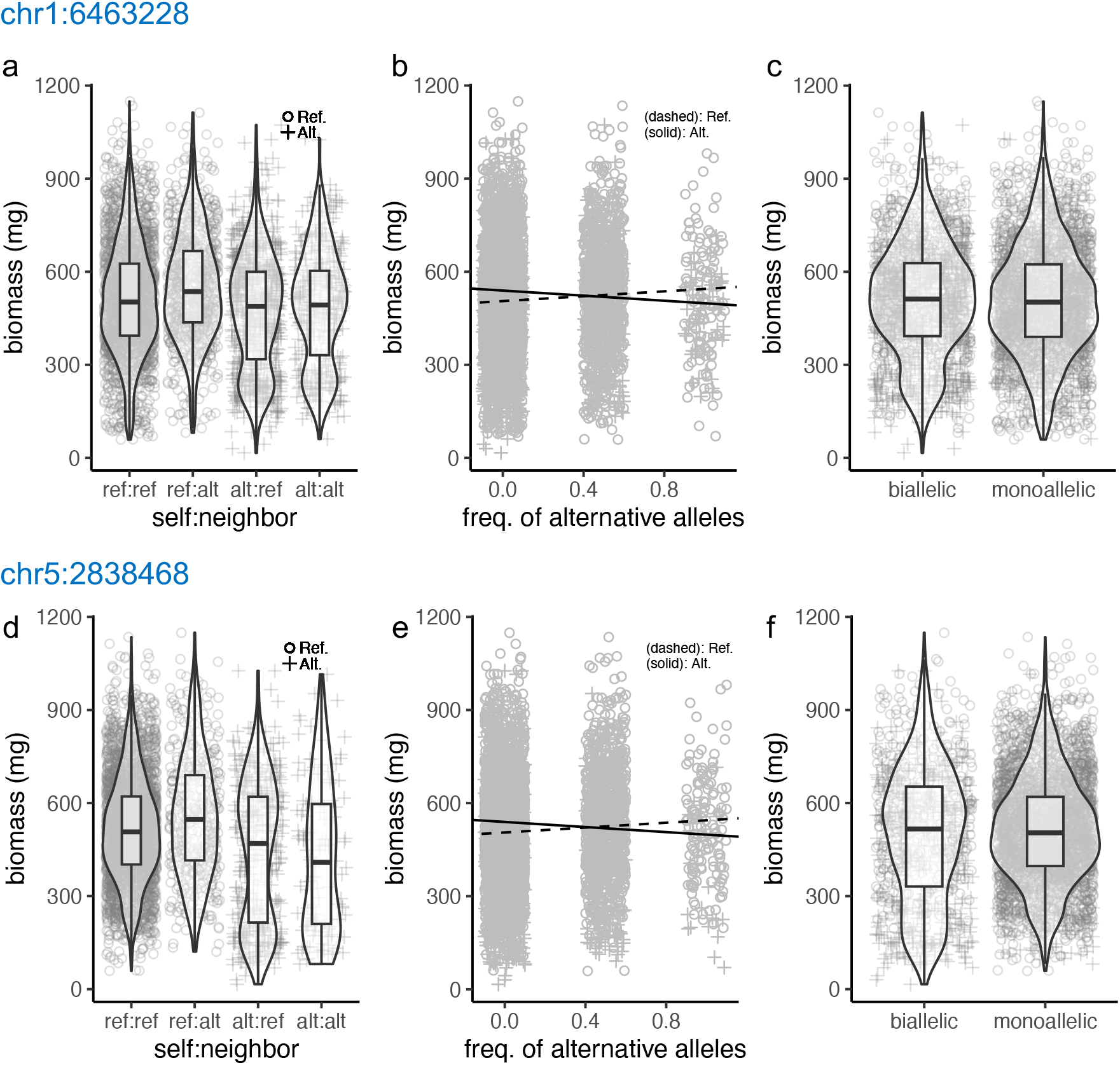
Patterns of individual biomass at two SNPs with significant *β*_s×n_ in GWAS. The upper **a-c** and lower **d-f** panels exemplify significant SNPs on the first chromosome (chr1:6463228) and the fifth chromosome (chr5:2838468). The individual biomass is plotted against self:neighbor allelic combinations between reference (ref) and alternative (alt) alleles (**a** and **d**); frequency of alternative alleles (**b** and **e**), and the biallelic or monoallelic conditions (**c** and **f**). Box plots show the median with upper and lower quartiles, with whiskers extending to 1.5 × interquartile range.

**Figure S4.**
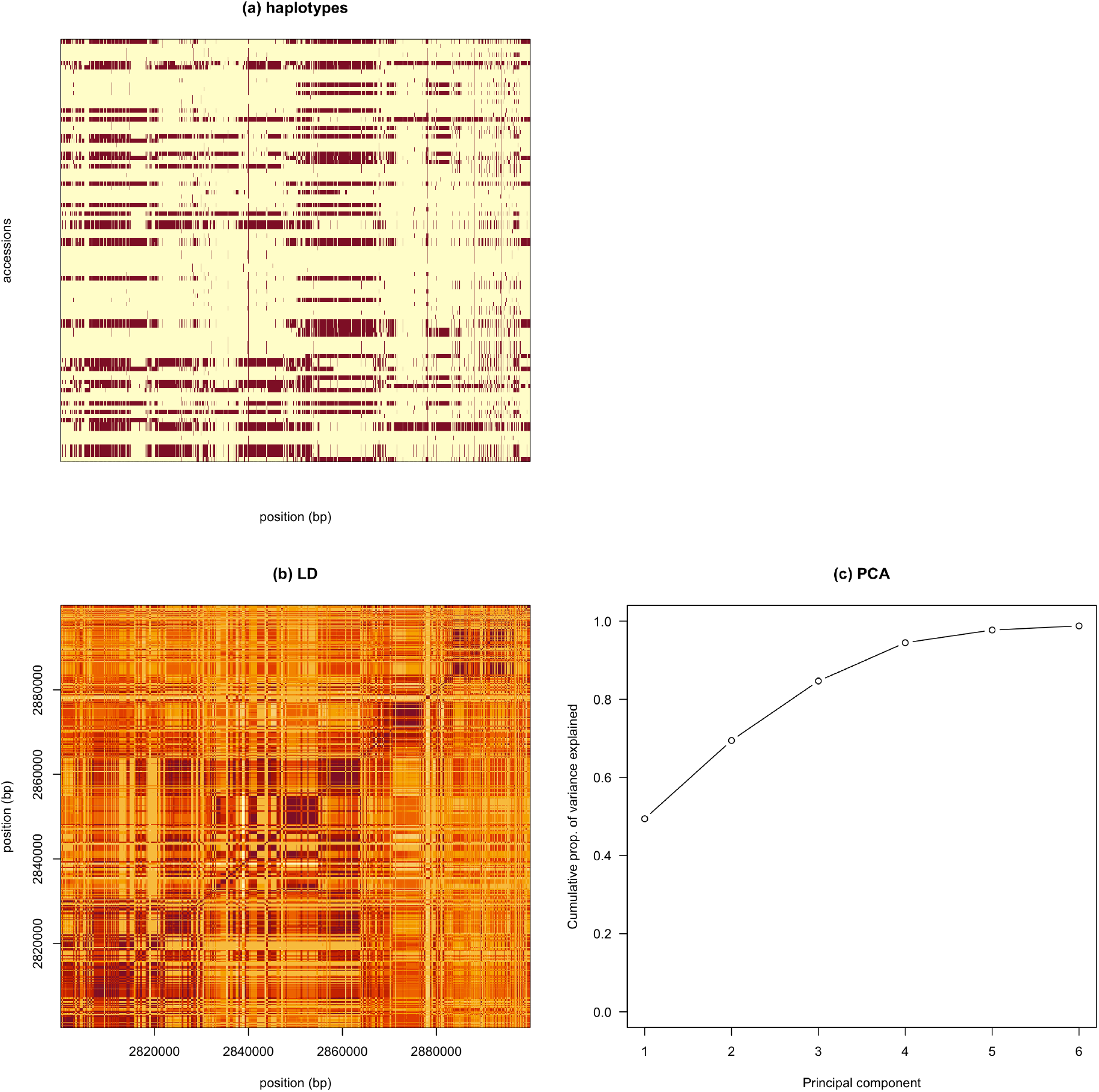
Patterns of haplotypes and linkage disequilibrium (LD) near the most significant SNP among 98 Arabidopsis thaliana accessions. Light and dark colors indicate reference (Col-0) and alternative alleles, respectively. (**a**) Haplotype patterns 100 kbp near the most significant SNP on the fifth chromosome. (**b**) Heatmap showing LD patterns in the same region as panel **a** among 98 accessions. (**c**) Principal component analysis (PCA) summarizing the LD pattern in the panel **b**. Cumulative contributions from the first to tenth principal components are shown.

**Figure S5.**
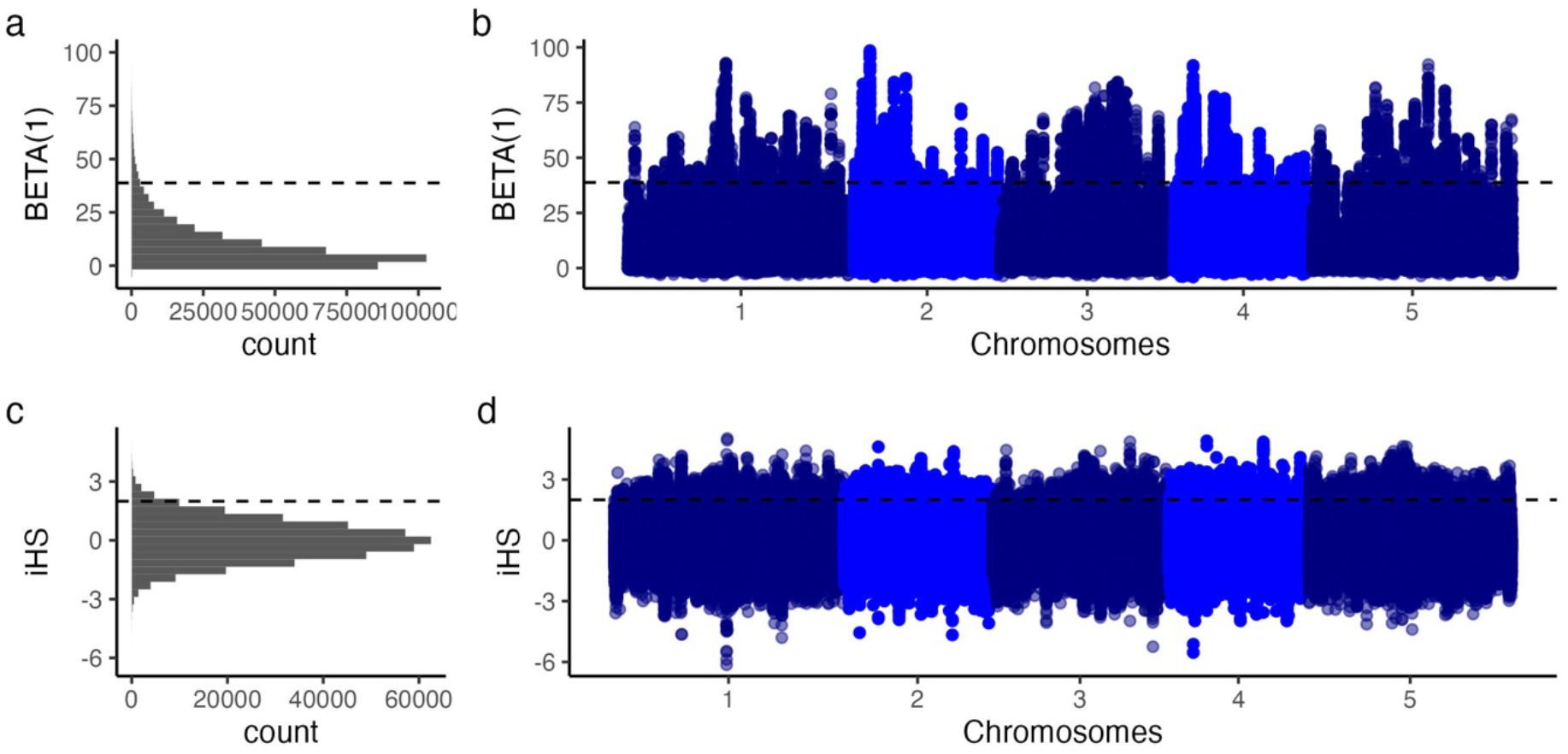
Genome-wide distribution of BETA (**a** and **b**) and iHS (**c** and **d**) statistics. Horizontal dashed lines indicate genome-wide 97.5 percentiles.

**Table S1.**
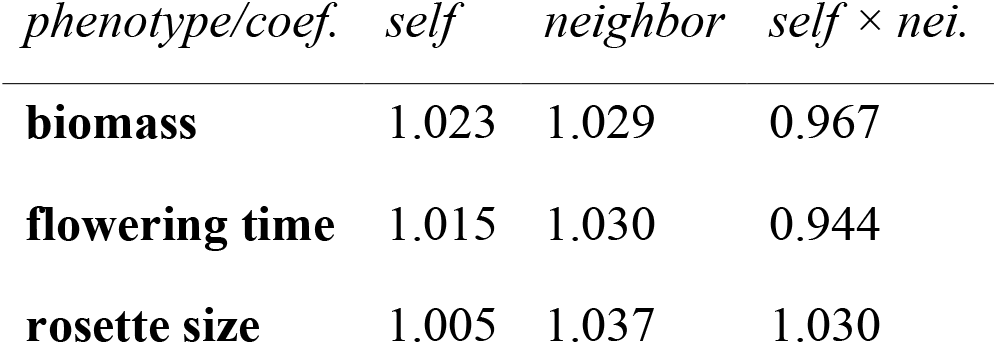
Genome inflation factors of GWAS *p*-values for the three phenotypes (biomass, flowering time, and rosette size) and coefficients (self, neighbor, and self-by-neighbor effects). The values of >>1 or <<1 indicate a genome-wide inflation and deflation of p-values, respectively.

| *phenotype/coef.* | *self* | *neighbor* | *self × nei.* |
| --- | --- | --- | --- |
| **biomass** | 1.023 | 1.029 | 0.967 |
| **flowering time** | 1.015 | 1.030 | 0.944 |
| **rosette size** | 1.005 | 1.037 | 1.030 |

